# Detection of Multidimensional Co-Exclusion Patterns in Microbial Communities

**DOI:** 10.1101/315473

**Authors:** Levent Albayrak, Kamil Khanipov, George Golovko, Yuriy Fofanov

## Abstract

**Motivation:** Identification of complex relationships among members of microbial communities is key to understand and control the microbiota. Co-exclusion is arguably one of the most important patterns reflecting microorganisms’ intolerance to each other’s presence. Knowing these relations opens an opportunity to manipulate microbiotas, personalize anti-microbial and probiotic treatments as well as guide microbiota transplantation. The co-exclusion pattern however, cannot be appropriately described by a linear function nor its strength be estimated using covariance or (negative) Pearson and Spearman correlation coefficients. This manuscript proposes a way to quantify the strength and evaluate the statistical significance of co-exclusion patterns between two, three or more variables describing a microbiota and allows one to extend analysis beyond microorganism abundance by including other microbiome associated measurements such as, pH, temperature etc., as well as estimate the expected numbers of false positive co-exclusion patterns in a co-exclusion network.

**Results:** The implemented computational pipeline (CoEx) tested against 2,380 microbial profiles (samples) from The Human Microbiome Project resulted in body-site specific pairwise co-exclusion patterns.

**Availability:** C++ source code for calculation of the score and p-value for 2, 3, and 4 dimensional co-exclusion patterns as well as source code and executable files for the CoEx pipeline are available at https://scsb.utmb.edu/labgroups/fofanov/co-exclusion_in_microbial_communities.asp

**Supplementary information:** Supplementary data are available at *biorxiv* online.

## 1 Introduction

Every life-supporting environment is colonized by microbes. Understanding the complex relationships among microorganisms opens new possibilities in controlling microbial communities (MC) and have a variety of applications from improving human health to reducing negative effects of microbial activities in industrial settings, such as microbiologically influenced corrosion, product spoilage, and formation of harmful biofilms.

Over the last decade, High Throughput Sequencing (HTS) technology has become a key component of a large variety of microbiome studies. Since whole genome sequencing (WGS) remains relatively expensive, and requires extensive computational resources to perform analysis on data which may consist of up to 95% host DNA, many of the recent studies have been focused on marker gene sequencing approach where selected genes (usually highly conserved such as 16S/18S rRNA or COI) are amplified, sequenced, and used to identify microorganism profiles of microbiota samples (Srivathsan *et al.*, 2015). The first steps of the analysis of this data include quality assessment of sequencing reads, clustering, taxonomic annotation, and normalization resulting in relative abundance profiles which may contain thousands of organisms identified across hundreds of samples. The downstream analysis of these profiles typically focuses on estimation of microbial diversity of individual samples and/or search for microbes exhibiting distinctive abundance patterns across specific groups of samples such as microorganisms highly abundant in one group and absent in another.

This approach has already resulted in discoveries of microorganisms potentially associated with colorectal cancer (Tjalsma *et al.*, 2012; Kostic *et al.*, 2012; Ahn *et al.*, 2013; Flanagan *et al.*, 2014; Sun and Kato, 2016), early gastric cancer (Lee *et al.*, 2016), prostate cancer (Perry and Lambert, 2011), and cystic fibrosis (Zhao *et al.*, 2014). Many recent studies (He *et al.*, 2017; Mainali *et al.*, 2017; Fisher and Mehta, 2014) have been dedicated to the identification of competitive, symbiotic, and commensal (Reshef *et al.*, 2011) relationships between members of MC, focusing on identification of microorganisms for which pairwise abundances can be modeled using linear, exponential, or periodic (for time series) functions.

Since manual search for even the simplest of patterns (such as Pearson or Spearman correlation (Pearson, 2006; Fieller *et al.*, 1957) in microbial abundance profiles is virtually impossible (the number of pairs of microorganism profiles to be inspected is proportional to the square of the number of microorganisms under consideration 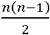, where n is the number of microorganisms in the dataset), various algorithms and computational pipelines have been developed to perform such tasks automatically. For example, several computational pipelines employ Pearson or Spearman correlation to identify the most significant positively and negatively correlated microorganism profiles (Barberán *et al.*, 2012; Cardinale *et al.*, 2015; Maruyama *et al.*, 2014; Wang *et al.*, 2016; Goodrich *et al.*, 2014; Jackson *et al.*, 2018). SparCC (Friedman and Alm, 2012) improves on correlation search by correcting undesirable effects of compositional data. Software packages such as CoNet (Faust *et al.*, 2012) employ several similarity measures such as Kullback–Leibler divergence (Kullback, 1959), Pearson correlation (Pearson, 2006) and Spearman correlation (Fieller *et al.*, 1957), as well as mutual information (MI) (Cover and Thomas, 2005) to identify the highest scoring pairwise relationships. Among these methods, only MI and MIC (Reshef *et al.*, 2011) algorithms are capable of identifying nonlinear and non-functional (impossible to represent in y=f(x) form) pairwise relations by using mutual information and maximum mutual information scores (see Weiss *et al*. (2016) for the review and performance comparison). These two methods, however, lack the ability to discriminate against intuitively difficult to interpret patterns (Figures 1a, b) and can miss some important relationships. For example, mutual exclusion among microorganisms (Figures 1c-f), produce low MI and MIC scores and as result such patterns will not appear on the top of the list of significant relationships identified and reported by the MIC pipeline (Reshef *et al.*, 2011). It is also obvious, that complex relationships among members of microbial communities can include two, three, and more organisms. To date however, no methods for identification of such multidimensional microbial appearance/abundance patterns have been adopted by the scientific community. In theory, MI and MIC approaches can be extended to identify patterns which include three, four, or more organisms but the computational complexity of such a task renders these methods impractical.

**Figure 1.**
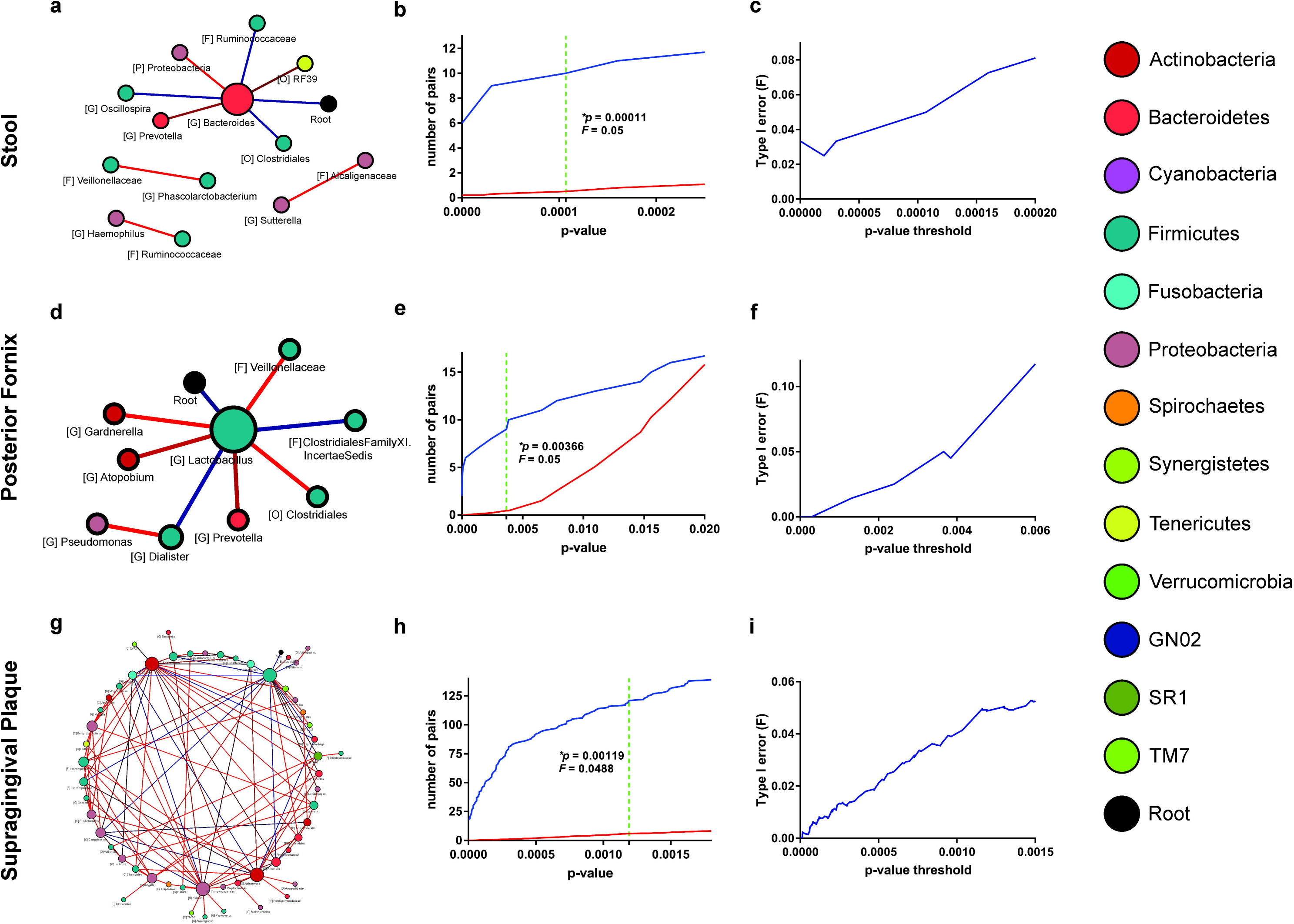
Non-random patterns (associations) including two and three organism profiles. a, b – intuitively difficult to interpret patterns with high MIC score; c – co-exclusion pattern involving 3 organisms; d, e, f – pairwise co-exclusion between three organisms involved in the 3-dimensional co-exclusion pattern. Values in the figure represent mutual information (MI); maximal information coefficient (MIC); Pearson correlation (ρ); co-exclusion coefficient (c).

*Co-exclusion* is arguably one of the most important patterns to be identified in microbial communities. Knowing which microorganisms are unable to tolerate each other’s presence or can replace one another in the community opens an opportunity to manipulate and control the microbiota, guide microbiota transplantation, and personalize the choice of microorganisms for probiotic treatments. Examples of recently discovered co-exclusion relationships include Wolbachia sp. co-exclusion with Asaia sp.(Hughes *et al.*, 2014) and Flaviviruses (Dengue, Zika) (Schnettler *et al.*, 2016; Bian *et al.*, 2010); and co-exclusion between Lactobacillus *spp*. and Fusobacteria *spp*. (Sobhani *et al.*, 2013), which are associated with elevated risk of colorectal cancer, may open new opportunities in cancer prevention.

It is important to emphasize however, that mutual exclusion/avoidance pattern is not anti-correlation (negative Pearson or Spearman correlation). The nature of co-exclusion does not depend on the abundance of given species but (in an ideal case) has to be based only on the fact that one microorganism can be present (in any abundance) only if the other microorganism is absent (and vise-versa). This type of relationship cannot be described in terms of a functional relationship (mathematically represented as a function) and requires a different equation type of representation which in the two dimensional case can be defined as: *X_a_ X_b_* = 0, where *X_a_* and *X_b_* are abundances of two organisms (a and b).

The goal of the presented study is to introduce a quantified definition of the strength and statistical significance of multidimensional co-exclusion patterns between variables describing microbial communities as well as estimate the correspondence between statistical significance threshold of individual co-exclusion patterns and expected numbers of false positive co-exclusion patterns in a co-exclusion network.

## 2 Methods

### 2.1 Goodness of fit for two-dimensional co-exclusion patterns

The coefficient of determination (*R*^2^) is traditionally used to quantify the goodness of fit of experimental observation data

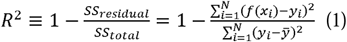

points {*y_i_*; *x_i_*} *i* = 1, …, *N* to the given functional (*y* = *f*(*x*)) model. In general terms, it reflects how much better the given model fits the experimental data in comparison with a base model and is mathematically defined as the ratio between the deviation of experimental observations from the model and the deviation from the “base model” which is usually chosen as the average (*y* = *y̅*):

Using the same principles and assuming that the co-exclusion pattern is described by the equation: *xy* = 0, the deviation of each experimental observation from the co-exclusion model can be defined as a geometric fit, which in this particular case is the shortest distance of the data point to the closest axis and *SS_residual_* can be defined as:

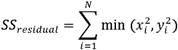

Since the deviation from the co-exclusion model is calculated using distances from both (x and y) axis, the base model also must include averages of both variables: {*x* = *x̅*; *y* = *y̅*} which allows definition of *SS_total_* as:

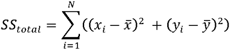

Thus the goodness of fit into the co-exclusion pattern *co-exclusion coefficient* (*c*) can be introduced as:

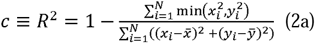

It is important to mention however, that in contrast with the goodness of fit for functional relationships (formula 1) the value of the *co-exclusion coefficient* (formula 2a) can be affected by linear transformation of one of the variables. In fact, its value can be artificially “improved” by multiplying all the values associated to one of the variables by a very small coefficient (*α*). In order to make the *co-exclusion coefficient* independent of linear transformations (which also allows calculation of its value for variables measured in different scales such as bacterial abundance vs. temperature, salinity, or pH), the linear transformation coefficient (*α*) must be included in the definition of *co-exclusion*

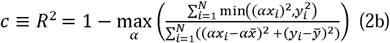

*coefficient:*

In this case (formula 2b), the *co-exclusion coefficient* is defined as the lowest (worst) *co-exclusion* value across all possible values of the linear transformation coefficients (Figures 2a, b).

**Figure 2.**
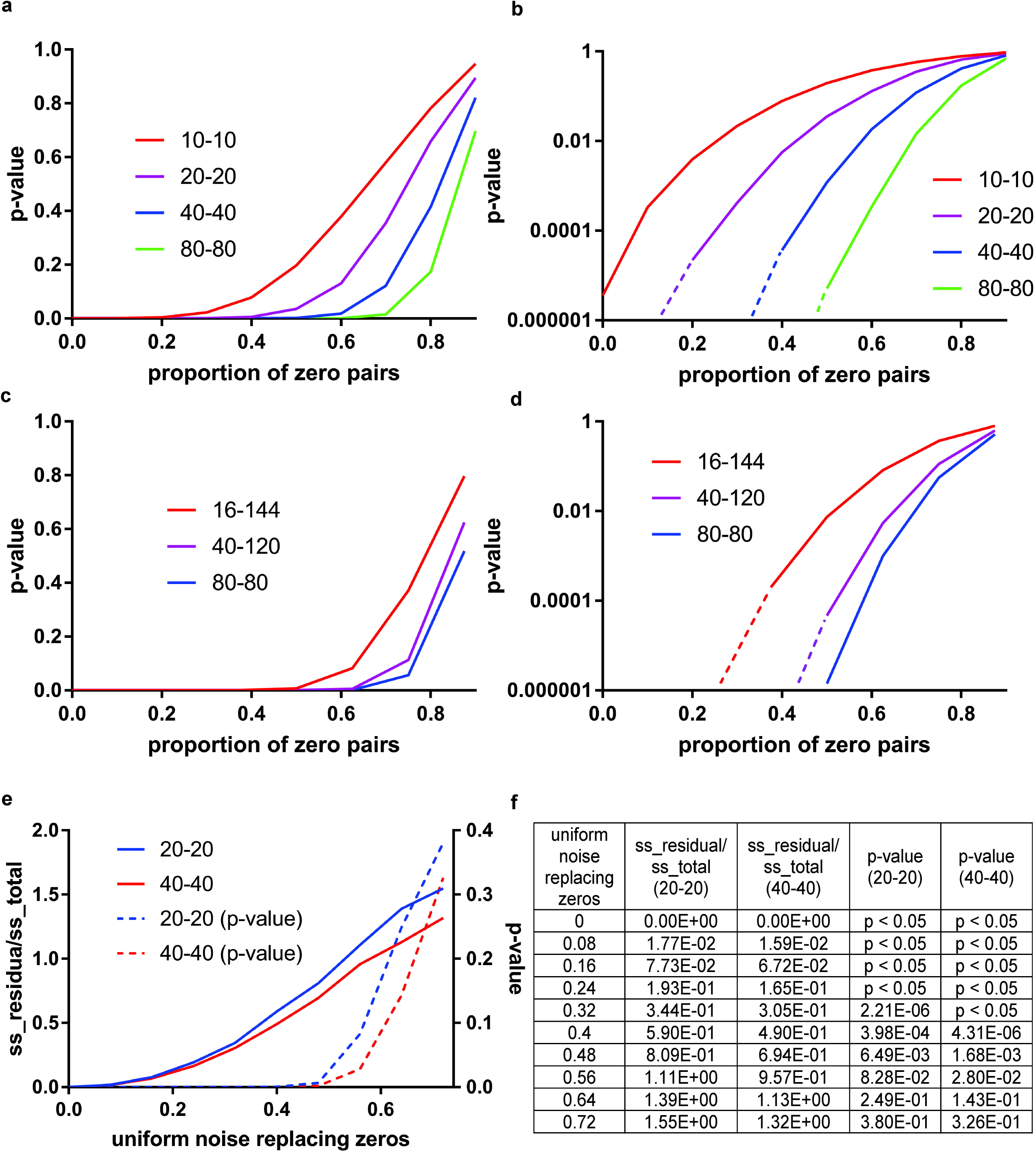
Effect of linear transformation on the goodness of fit for two dimensional (a-b) and three dimensional Type-1 (c, d) and three dimensional Type-2 (e, f) co-exclusion. Figures b, d, and f show the effect of linear transformation on the 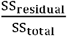 ratio (formulas 2b, 3a, and 3b).

### 2.2 Statistical significance of the co-exclusion pattern and the data properties influencing it

Regardless of how strong the value of the co-exclusion coefficient is, certain properties of the data can significantly affect its ability to reflect what is intuitively considered as a mutually exclusive relationship. The most striking example of such effects would be if the co-exclusion coefficient is calculated between two variables out of which one is always zero. In this case all the observations will be located on one axis resulting in a “perfect” co-exclusion score. Another case where false but high scoring co-exclusion patterns can be observed simply by chance is when both variables have only a few non-zero values across a large number of observations and therefore, the data becomes dominated by a large number of zero-pairs (both variables equal to zero) which perfectly fit in the co-exclusion pattern (Figure 2a). In fact, in zero-pair rich datasets, high scoring but false co-exclusion patterns can appear between pairs of independent variables due to the low probability of both variables having non-zero values in same sample (observation).

Since no assumption regarding the distributions of the values of the variables can be made, the statistical significance (such as p-value) of the observed co-exclusion pattern can be estimated using a *bootstrapping* approach. In this case, p-value can be calculated as a ratio of the number of the simulated bootstrap attempts in which the co-exclusion coefficient values are better or equal to the co-exclusion score of the original data over the total number of bootstrap attempts. For all the results presented in this manuscript, we have used up to 100,000 bootstraps for each pair of variables. It is important to emphasize however, that in order to improve the computational performance, several additional criteria have been used to discontinue the bootstrapping iterations when it becomes clear that the expected p-value will be higher than the predefined threshold.

To date, several authors have raised concerns over the significant effects of presence of zero-pairs (observations where both variables have zero or very low values) on results of correlation analysis between OTU profiles (Xu *et al*., 2015; Kaul *et al.*, 2017; Weiss *et al.*, 2017). To explore these effects on co-exclusion patterns, we have performed several computational experiments using simulated data. For four sample sizes (20, 40, 80, and 160), we have randomly generated 10 datasets with perfect co-exclusion patterns where all non-zero values were generated randomly in the 0.2-0.9 interval and were paired with zero values for the corresponding variable. The proportion of non-zero values on each axis were equal (50%-50%). During the sequential steps of the simulation 0%, 10%, 20%, …, and 90% of the data points were replaced by zero-pairs. Not surprisingly (Figures 3a,b), the p-values of the observed co-avoidance pattern improve when number of samples increases, while increasing the proportion of zero-pairs has the opposite effect. This observation highlights the connection between the number of samples and proportion of zero-pairs with the expected co-exclusion pattern’s p-value. For example, for sample size 80 and 50% of zero-pairs, the p-value of the perfect co-avoidance pattern is expected to be equal or less than 10^−5^.

**Figure 3.**
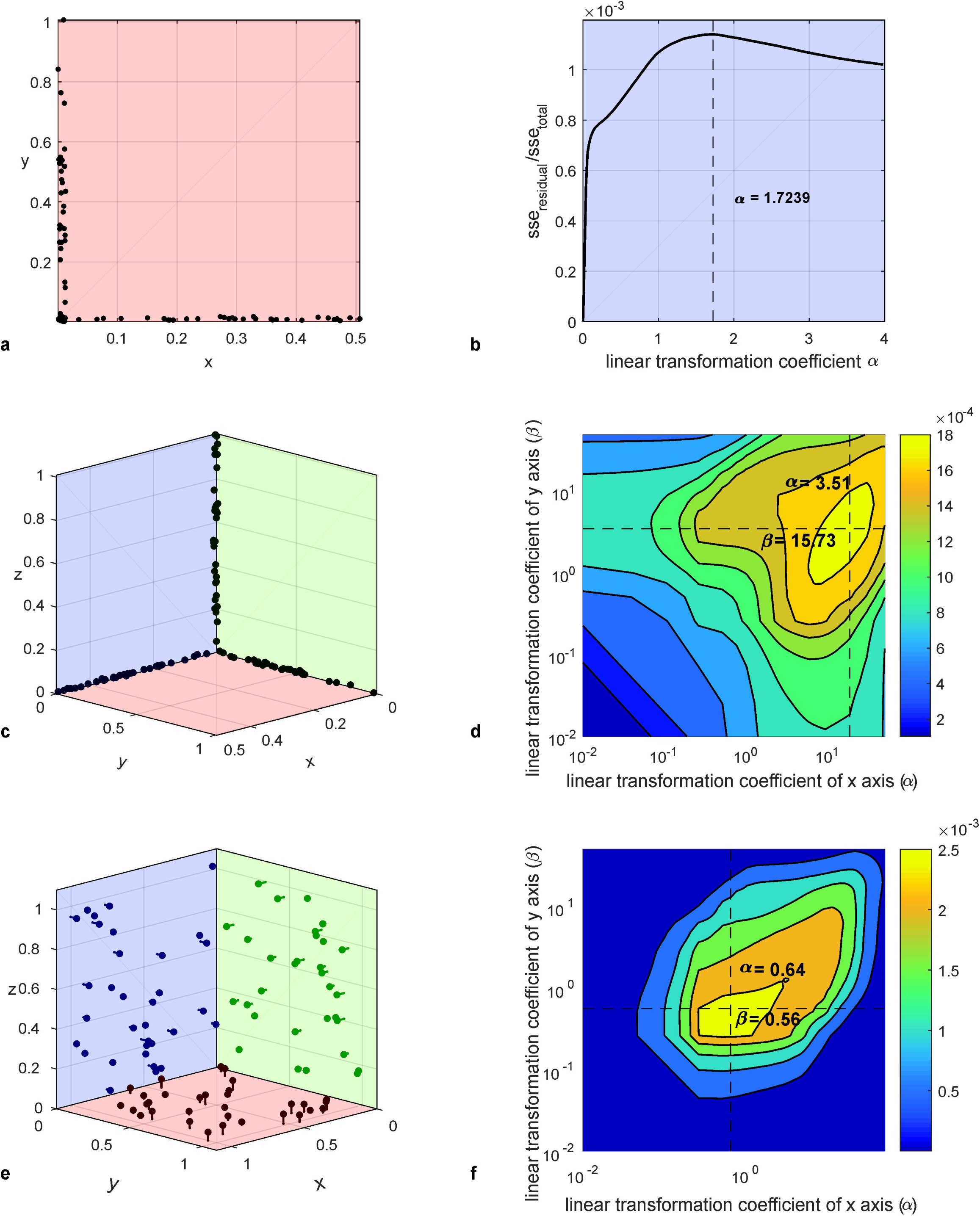
(a) The effects of zero-pairs and sample size on the p-value on the “perfect” co-avoidance pattern; (c) the effects disproportional presence of no-zero values between two variables zero-pairs on the “perfect” co-avoidance pattern; (b) and (d), same data in log scale; (e) and (f), Effects of zero-pairs and noise on the co-exclusion (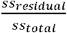) score.

The disproportional presence of non-zero values between two variables can also affect the p-value of a perfect co-exclusion pattern. The simulation results on Figures 3c,d show that while such an effect is minimal for 25% to 75%, it becomes stronger for 10% to 90% ratio, especially for cases with high number of zero-pairs. Random noise and measurement errors can also affect p-values of perfect co-exclusion patterns. It is important to mention however that introduction of low level (<10%) noise (Figures 3e,f) affects the value of the co-exclusion coefficient, but not its significance (p-value).

### 2.3 Goodness of fit for multidimensional co-exclusion patterns

There are two different ways how co-exclusion can be defined

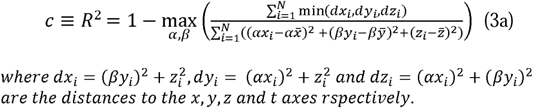

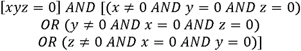

in the case of three variables (3D co-exclusion). The Type-1 co-exclusion characterizes the situation when each organism avoids every other organisms in a group (Figures 2c,d). In this case, the co-exclusion pattern is represented by three axes and geometric distance for each experimental observation is calculated as the distance to the closest axis and the co-exclusion model has to be represented as:

Similar to formula 2b, to exclude the dependence of the co-exclusion score from linear transformations, 3 dimensional Type-1 co-exclusion definition will need to include two transformation coefficients (α and β). The Type-1 3D co-exclusion score can be defined as:

Not surprisingly, to express Type-1 three-dimensional co-exclusion, pairwise co-exclusion patterns have to be present between each pair of variables involved in the 3D pattern. This allows one to establish simple exclusion rules and reduce the computational complexity of the search for multidimensional co-exclusion patterns from *O*(*m*^3^) to *O*(*m*^2^), where m is number of variables considered in the pattern:

Rule 1. Two variables exhibiting two dimensional co-exclusion pattern with zero or small number of observations where both organisms are absent in the samples cannot be part of any higher dimensional co-exclusion pattern;
Rule 2. Only variables exhibiting all three pairwise co-exclusion patterns simultaneously can form a 3D Type-1 co-exclusion pattern.

Another (Type-2) three-dimensional co-exclusion pattern could be defined as the case when each pair of organisms could co-exist, but all three cannot be present simultaneously (Figures 2e-g). In this case, the co-exclusion model can be represented as three axis planes: x=0 (yz plane), y=0 (xz plane), and z=0 (xy plane) and the co-exclusion coefficient can be calculated using minimal distance from these planes and the co-exclusion model has to be represented

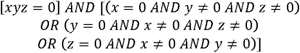

as:

The Type-2 3D co-exclusion score can be defined as:

Interestingly, in this definition Type-2 co-exclusion pattern will show high co-exclusion score if variables are exhibiting Type-1 co-exclusion, but not vise-versa. A simple (but less effective) exclusion criterion can be used to avoid unnecessary calculations

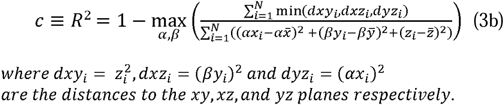

to identify if any three variables can be involved in Type-2 3D co-exclusion: every two variables must have a significant number of zero-pairs: observations where one or both organisms are absent.

Co-exclusion patterns can be defined for higher number of variables (dimensions). For example, co-exclusion in four

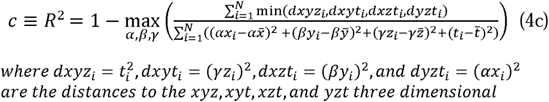

dimensions can be calculated using distances from axes (formula 4a), axis planes (formula 4b), as well as three dimensional subspaces (formula 4c) where t stands for the fourth dimension (fourth variable):

While formulas 2b, 3a,b and 4a-c can provide a numerical value for the strength of the co-exclusion, the above described bootstrapping approach can be used to estimate its confidence.

### 2.4 Co-exclusion network: Type I/II errors and expected fraction of false positive pairs

Considering the large number of combinations of two, three, and more variables tested independently in search for co-exclusion patterns, a substantial number of statistically significant patterns can be observed simply by chance (multiple hypothesis testing problem). Assuming that the p-values of individual co-exclusion patterns reflect the probability of these patterns to appear by chance (false positive), one can use a bootstrapping approach to identify the correspondence between p-value thresholds and expected number/fraction of encountered false positive co-iterations (Figures 4b, 4e, and 4h) allows one to estimate the expected fraction of the false positive edges (Type I errors) in the co-exclusion network (Figures 4c, 4f, and 4i) as a function of the p-value threshold. In all the microbiota datasets presented in the results section we have generated 20 randomized (shuffled) datasets. All the networks (Figure 4 and Supplementary Materials document, section 2) were generated using 0.05 threshold for the expected fraction of false positive pairs (5% false positive rate).

**Figure 4.**
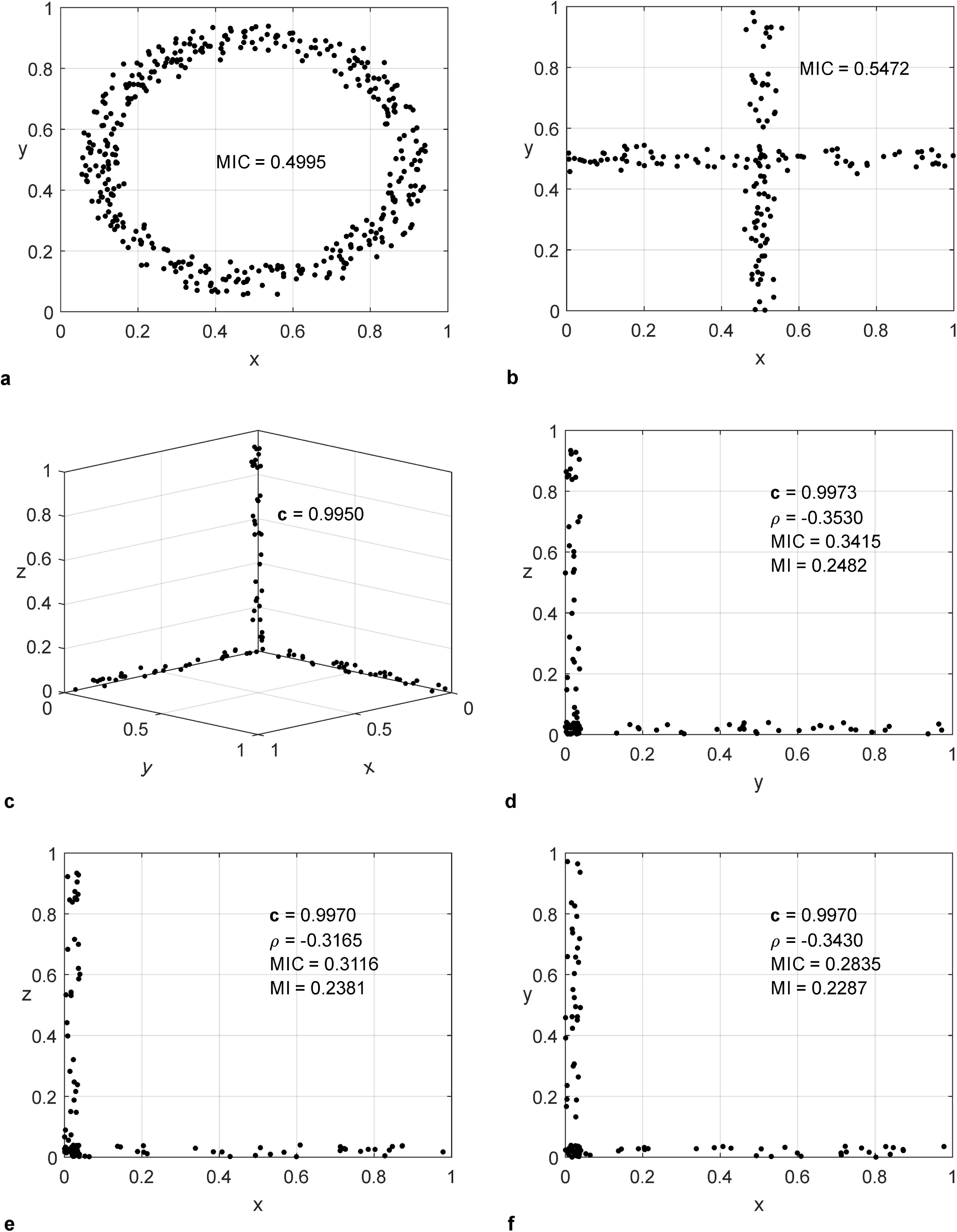
Examples of co-exclusion networks and p-value distributions using Stool, Posterior Fornix, and Supragingival Plaque data sets from the HMP. The co-exclusion networks (a, d, and g) are created using 5% cutoff for the allowed fraction of false positive pairs (F). Strong co-exclusion (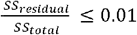) is indicated in bright red edges with a transition to blue as the co-exclusion score approaches 0.2. Node colors reflect different taxonomy assignments at Phylum level and node sizes are proportional to the number of co-exclusion relationships. The cumulative distributions of the p-values for pairs in observed (blue) and simulated (red) networks (b, e, and h) are shown with a green dashed line which represents the p-value threshold (^*^p) used to create the networks. The expected fraction of the false positive edges (Type I errors) in the co-exclusion network as a function of the pairwise p-value threshold is shown in the 3^rd^ column (c, f, and i).

The estimation of the Type II error (false negative) in reconstruction of any network describing microbial interactions is quite challenging. In general, it requires the presence of a model describing the ground-truth of the relationship between variables using artificial assumptions. For example, in the Friedman and Alm manuscript (Friedman and Alm, 2012), the false negative edges in the network created using SparCC algorithm were defined as edges present in the SparCC generated network but absent in the network created using Pearson correlation. It is important to emphasize that in the presented co-exclusion network reconstruction process, the selection of any cutoff for pairwise co-exclusion patterns.

In this approach, the identification of co-exclusion patterns in the original data is supplemented by several iterations of identification of co-exclusion patterns in randomized datasets (Null Model) generated by shuffling and re-normalization of the original dataset. Comparison of the distributions of the p-values in the original and p-values of simulated data averaged across all exclusion p-values results in the rejection of some number of “perfectly fine” co-exclusion patterns which cannot be estimated, at least not without including additional assumptions/model(s) in consideration.

### 2.5 Implementation

The C++ source code for the calculations of the score and p-values for 2, 3, and 4 dimensional co-exclusion patterns as well as source code and executable for the computational pipeline (CoEx) for the reconstruction of 2-dimmentional co-exclusion networks is available at https://scsb.utmb.edu/labgroups/fofanov/co-exclusion_in_microbial_communities.asp).

The presented computational pipeline also performs bootstrapping simulations for (a) each pairwise co-exclusion pattern and (b) the resulting network in a single run. Each run of the pipeline results in two tables (text files): one containing information (such as co-exclusion score and p-value) on co-exclusion patterns between pairs of OTUs within the original data and one containing the simulated network p-value distribution results (more details are available in the Supplementary Materials document). The pipeline source code can be compiled using g++ for Linux or Microsoft Visual Studio C++ compilers. The software is single-threaded and requires minimum 8GB memory. The run time can vary depending on the resulting number of pairwise relationships as intensive bootstrapping must be completed for each reported pair. For example, the exhaustive analysis of 4,851 OTU pairs for identification of statistically significant co-exclusion patterns using 160 stool samples involving 99 OTUs shown in Figure 4 took less than ~18 minutes to complete on an Intel^®^ Core^™^ i7-3630QM CPU @ 2.40GHz ASUS laptop with 12.0 GB installed RAM (100,000 trials per pair and 20 network replicates).

The co-exclusion network (Figure 4) has been visualized using Cytoscape 3.4.0 (Shannon *et al.*, 2003) visualization platform using the yFiles Organic network layout algorithm. MIC calculations (Figure 1) have been performed in MATLAB (www.mathworks.com/products/matlab) using minepy (Albanese *et al.*, 2013).

### 2.6 Human Microbiome Project data

The data used in the analysis is originated from The NIH Human Microbiome Project (Peterson *et al.*, 2009) and contained 18 datasets associated with 16 body sites. Microbial profiles for 2,910 samples have been downloaded from the project website as of December 2016 in text format (HMQCP - QIIME Community Profiling v13 OTU table). Samples representing significantly low (less than 2,000) and significantly high (over 50,000) number of sequencing reads were excluded from the analysis. The microbial profiles of the remaining 2,380 samples, vary from 67 for Posterior Fornix to 200 for Antecubital Fossa, have been normalized against the total number of reads in each sample and transformed into relative abundance profiles merged to Genus taxonomy level for each body site resulting in 619 profiles. Analysis has been performed for each body site individually. For each body site under consideration, microorganism profiles present in less than 5% of samples have been excluded from the analysis.

## 3 Results: Co-exclusion networks reconstruction

To evaluate ability of the proposed approach to identify co-exclusion patterns, we have applied the developed pipeline (CoEx) to the microbial profiles obtained from the Human Microbiome Project. The list of the identified co-exclusion patterns, results of the bootstrapping based false positive edge estimations, as well as the networks figures are available in the Supplementary Materials document.

Interestingly, no correlation has been observed between the number of organisms present in the dataset and the number of detected co-exclusion patterns, or the number of organisms involved in co-exclusion relationships. The highest fraction of organisms involved in co-exclusion patterns were identified in samples associated with the oral cavity: Tongue Dorsum (77.78%), Palatine Tonsils (64.65%), Supragingival Plaque (62.22%), Subgingival Plaque (61.70%), and Throat (46.96%, see Supplementary Materials document for more details), which could suggest that this environment can support a large variety of different (organism profiles) microbiomes. In Saliva samples, a significantly smaller fraction of organisms involved in co-exclusion relationships was identified (20.35%) when compared to the Throat dataset. Both data sets consist of similar number of samples and organisms (130 samples and 113 OTUs and 140 samples and 115 OTUs for Saliva and Throat data sets respectively). This could be due to the specific properties of Saliva (chemical/food/dead epitalia cell concentration, pH, etc.) and its wide spatial reach which can be seen as a temporary reservoir of detached members of the established microbiota present in the oral cavity.

The lowest fraction of the organisms involved in co-exclusion was found in Anterior Nares (1.55%) followed by Retroauricular Crease (3.70%) – despite large number of organisms found in Anterior Nares samples (200, largest number across all 16 body sites). This result however is not surprising considering that most of the microorganisms identified in these two sample types could be of environmental origin and do not represent the native “core” microbiota of this body site. In comparison to the oral cavity samples, a slightly lower fraction of organisms has been found in vaginal samples: Posterior Fornix (14.93%), Vaginal Introitus (14.58%), and Mid Vagina (11.76%).

It is also interesting to mention that not all the detected highly significant patterns appear to have a good co-exclusion score. For example, the weak co-exclusion coefficient score (0.192253) between Dialister (Genus) and Lactobacillus (Genus) in the Posterior Fornix network (Figure 4d and Supplementary Materials document, section 2.2) has been found to have a very low p-value (0.000289).

## 4 Discussion

Co-exclusion is one of many non-random association patterns which can be detected in microbial communities. This pattern however, is key to understanding complex relations in microbial communities. Knowing which organisms are incompatible, or can simply replace each other in MC can guide the development of patient-specific probiotics and improve microbial transplantation strategies.

However, it is important to keep in mind that strong and statistically significant co-exclusion does not necessarily reflect competition or inability of organisms to co-exist in the same microbial community. Artificial co-exclusion (especially on the OTU level) may appear as an artifact of the taxonomical annotation. Moreover, in some datasets, co-exclusion can be observed simply because microbial compositions originated from very different environments. For example, by analyzing posterior fornix and gut (stool) microbiota profiles together, one can observe co-exclusion pattern between organisms specific (uniquely present) in each of these two environments. Specifically, Clostridium (from family Erysipelotrichaceae), Subdoligranulum, and Akkermansia absent in posterior fornix samples and present in 81%, 78%, and 52% of stool samples will exhibit co-exclusion patterns with Finegoldia, Ureaplasma, and Herbaspirillum absent in stool samples and present in posterior fornix (39%, 31%, and 38%).

It is also necessary to keep in mind that a high co-exclusion score with low statistical confidence cannot be interpreted as absence of co-exclusion. It simply reflects a lack of statistical evidence (not enough data points) to confirm the presence of co-exclusion.

Surprisingly, in some of the analyzed microbial communities several high confidence (based on p-value) co-exclusion patterns have been found to have relatively low co-exclusion scores. This observation makes us hypothesize that to a certain extent, the pairs of variables showing high confidence, but low co-exclusion scores may represent some type of co-presence, but we would like to keep this discussion outside the scope of this manuscript.

The overall complexity of the pairwise co-exclusion networks can reflect the diversity of microbial communities which are able to populate a given environment. Traditional directed/undirected graph representation however, cannot be used for multidimensional (except maybe Type-1) co-exclusion patterns. The ways how these and many other types of multidimensional relations can be visualized require development of novel approaches. It is also important to keep in mind that the best way to characterize the functional activity of microbiomes would be observing complete profiles of present (DNA) and expressed (RNA) genes, which at this time is cost prohibitive. The presented approach, however, is expected to be applicable to gene profile data when available.

## Acknowledgements

The authors would like to thank Iryna Pinchuk and Grant Hughes for the fruitful discussions and constructive comments.

